# Accelerated High-throughput Plant Imaging and Phenotyping System

**DOI:** 10.1101/2022.09.28.509964

**Authors:** Talha Kose, Tiago F. Lins, Jessie Wang, Anna M. O’Brien, David Sinton, Megan E. Frederickson

## Abstract

The complex web of interactions in biological communities is an area of study that requires large multifactorial experiments with sufficient statistical power. The use of automated tools can reduce the time and labor associated with experiment setup, data collection, and analysis in experiments aimed at untangling these webs. Here we demonstrate tools for high-throughput experimentation (HTE) in duckweeds, small aquatic plants that are amenable to autonomous experimental preparation and image-based phenotyping. We showcase the abilities of our HTE system in a study with 6,000 experimental units grown across 1,000 different nutrient environments. The use of our automated tools facilitated the collection and analysis of time-resolved growth data, which revealed finer dynamics of plant-microbe interactions across environmental gradients. Altogether, our HTE system can run experiments of up to 11,520 experimental units and can be adapted to studies with other small organisms.

## Introduction

The size of the standard scientific dataset has consistently increased through time, as larger datasets allow detection of more or more-complex effects. However, data demand has recently skyrocketed with the rise of artificial intelligence (AI) and machine learning (ML) techniques. AI and ML have transformed inference, but their utility is only realized when the training databases are expansive. High-throughput experimentation (HTE) is the key to creating large databases for AI and ML (Eyke et al., 2021). The primary challenge in HTE is the substantial time and cost to adapt existing experimentation methods to novel automated systems, which can dissuade researchers. Yet, pre-automation methods generally depend on intensive manual labor, creating a bottleneck and long-term higher costs to achieve high-throughput. Therefore, functional and cost-friendly HTE systems that are compatible with existing experimental methods and adaptable across experimental systems can substantially advance scientific research, especially in the biosciences (Kehe et al., 2019).

The study of biology is often concerned with the interactive effects of one or more type of organism assembled in communities across differing environments. Yet, when considering interactive effects of multiple classes of organisms (Song et al., 2020), multiple environmental contexts (Birk et al., 2020; Y. Yang et al., 2020), or both (Shears & Ross, 2010), the scale of experiment required for inference exponentially increases. HTE tools that facilitate the setup of experiments can provide transformative results (Kehe et al., 2019). Likewise, at the end of the experiment, phenotyping and collecting data from each experimental unit is often a rate-limiting step (W. Yang et al., 2020). In developing HTE, biologists often phenotype via automated imaging and automated image processing (Vasseur et al., 2018; Zhang & Zhang, 2018). In HTE platforms, collecting phenotype data quickly, yet also with high accuracy, is critical to overcoming the phenotyping bottleneck. Error detection and elimination during image data processing helps reach the desired accuracy (Cao et al., 2019). Scientists often leverage computing languages such as C++, Python, or MATLAB to create feature extraction algorithms for the phenotype of interest, such as by incorporating color thresholding for detecting plant tissues (Choudhury et al., 2019) (Cox et al., 2021) (Zhang & Zhang, 2018).

Understanding host-microbiome interactions presents one biological subfield with a growing issue of experimental scale (Kehe et al., 2019). We know species interactions can vary across environments (Chamberlain et al., 2014), including host-microbiome interactions (David et al., 2020). Yet, the environment can vary in myriad ways, presenting a real challenge to biologists. Researchers often want to measure host phenotypes, host fitness, or turnover in microbiome composition across several environmental variables shifting together and individually, but they are limited to smaller-scale experiments (Ge et al., 2022).

Rapid growth and utility to environmental applications are two significant factors researchers must consider when selecting a study system in the field of host-microbiome interactions. Duckweeds hold promise as a study system (Acosta et al., 2021) (Jones et al., 2021). Duckweeds grow rapidly across a wide range of environmental conditions (Lasfar et al., 2007), making them well-suited for studying responses of host-microbiome interactions to environmental variation (O’Brien et al., 2020), as well as for developing applications such as bioremediation and nutrient recovery from wastewater, biofuel, and nutrition for animals and even humans (Bhanthumnavin & McGarry, 1971) (Caicedo et al., 2000) (Appenroth et al., 2018) (O’Brien et al., 2022).

Duckweeds, including our focal species *Lemna minor*, have leaf-like fronds that float on the water surface. Mother fronds produce daughter fronds from meristems on either side, which may remain attached even after producing grand-daughter fronds, resulting in frond clusters (Newton, 1977). The diameter of individual frond clusters varies from 1-15 mm. Handling many of these small floating duckweeds on liquid surfaces at the scale required for an HTE can easily overwhelm even experienced researchers, creating a major bottleneck. One method uses a special coating on inoculation loops to pick and move duckweeds, but changes the duckweed medium from liquid to solid (Jones et al., 2021). New approaches such as filtration systems (Gregori et al., 2017) can be built and employed for duckweed handling. However, these solutions are insufficient for HTE with duckweeds due to incompatibility with existing labware, which would require intermediate systems to interface new methods with existing setups.

Imaging of multiple duckweeds with a high-resolution system is practical due to their small features. However, image-phenotyping duckweed experiments are still far from being HTEs: (Caicedo et al., 2000) reports only 10 treatments with three replicates. Cox et al. studied duckweed growth using 12-well plates. The highest capacity duckweed experiment currently reported is 3,000 units in (Nguyen et al., 2018), which included neither imaging nor phenotyping. Reliable and consistent sample imaging plays a crucial role: only replicable imaging conditions yield the level of image standardization required for batch processing (Zhang & Zhang, 2018), yet these have not been realized at the HTE scale for duckweeds. Automated imaging systems where duckweeds can be maintained inside a standardized imaging platform (as reported in, Cox et al., 2021), should provide the path forward.

Here, we demonstrate two automated systems focusing on accelerating experiment preparations and autonomous image-based phenotyping, which together allowed duckweed experiments to reach the HTE level. Our customized duckweed loading system integrates a liquid handling robot to ease labor-intensive duckweed handling in experiment preparation. Our automated imaging system takes images (or even videos) of samples throughout the experiment without interrupting or changing the experiment conditions, while our custom-built user interface runs a color thresholding algorithm to enable fast, manual-labor-free phenotyping of samples in image-processing. When combined, these three HTE platforms are capable of running experiments with up to 11,520 duckweed units. Most parts rely on automated processes and require only little supervision. In order to demonstrate our system capabilities, we designed an experiment with 6,000 duckweed units. Although we focus on duckweed experimentation in this paper, each system can be adapted to a variety of studies with small organisms.

## Materials and Method

### Experiment Design

We conducted an experiment investigating the effects of nutrient stressors on duckweed growth over time. Ten concentrations each of NaNO_3_, Ca(H_2_PO_4_)_2_, and KCl were crossed to generate a total of 1,000 different combinations of nitrogen (N), phosphorus (P), and potassium (K) levels. These combinations were crossed with two microbe treatments, present (inoculation with microbes isolated from plant line) or absent (inoculation with sterile liquid culture media). We then replicated these treatments across one genotype of the duckweed *Lemna minor* to reach 6,000 units. Each experimental unit was a tiny microcosm in one well of a 96-well plate (Sarstedt, 83.3925), containing duckweeds, and one combined treatment of N, P, K, and microbe presence. Each well received 240 µL of liquid growth media and nutrients followed by either 5 µL of microbial inoculum (20,000 cells/µL) or 5 µL of sterile culture media. Microbe treatments were blocked by plate to prevent cross-contamination while nutrient combinations were randomized across wells. Plates were sealed with a gas-permeable membrane (Sigma-Aldrich Z763624) to prevent contamination and reduce the evaporation rate of well solutions. Plates were monitored for 10 days in an environmentally controlled growth chamber at 22°C and 150 µmol^2^ lighting for 16 hours, and 18°C and dark for 8 hours. A total of 64 well plates were required to accommodate the high number of samples.

### Preparation of Duckweed Samples

Duckweed samples were originally collected from Cedarvale Pond (43°41’23.0”N 79°25’10.0”W) in Toronto, Ontario, and were clonally propagated from a single frond unit. Microbes were collected from crushed plant tissue and incubated on yeast mannitol agar (YMA) plates at 30°C for two days followed by storage at 4°C. Following the isolation of microbes, plants were sterilized by treatment with 1% NaOCl for 60 s. The clonal, sterilized line was maintained in sterile growth media (recipe adapted from (Krajnčič et al., 1995)) in glass jars.

The biggest challenge of a HTE duckweed experiment is the placement of duckweed samples into the wells. It requires tedious attention to pick up duckweed samples from a source jar since they are floating on the liquid surface freely. The small size of duckweed samples–typically 1-2 mm in diameter–increases the difficulty of such an operation. An experienced researcher can prepare a 96-well plate filled with duckweeds and other experimental components in 2-3 hours, but the job is highly repetitive. An automated system would reduce the manual labor. Hence, we set out to deliver one such automated system for this experiment.

### Autonomous Duckweed Loading System

We utilized an Opentrons-2 (OT-2) liquid handling system (Opentrons, OT-2 lab robot) to load the wells with duckweeds. This system was already able to fill well plates with liquids in custom experimental designs, and has a gantry system appropriate to handle duckweed samples. The OT-2 system was also integrated with a HEPA filter provided by the manufacturer in order to create a more controlled air-flow and hence sustain a sterile environment during the liquid handling or duckweed loading.

However, there aren’t suitable tips in the OT-2 system for the autonomous handling of duckweed. Typically, disposable plastic inoculation loops (Sigma Aldrich, Inoculating Needle/Loops, HS81121A) are preferred to catch the duckweed from a source jar. Hence, our goal was to design inoculation loop tips compatible with the OT-2 that can pick up and move duckweeds. BioPlas inoculation loop tips (BioPlas, 1 µL Astral inoculation loop, 7000) were able to pick up duckweeds from source samples, and have the added benefit of sterile packaging, allowing sterile operations between the wells. These loops are designed to be paired with the Astral Inoculation Loop Handles. However, the handles were not compatible with OT-2 pipette heads. We therefore modified the inoculation loops with 10 µL pipette tips so that OT-2 P300 pipette heads can engage with the inoculation loops and function as desired. The details regarding this modification can be found in the SI.

In addition to the 96-well plate to be filled with individual experimental duckweeds, a 12-well plate was employed as the source of all experimental duckweeds. In this source plate, wells were filled with DI water and packed with duckweeds. The density of the duckweeds and liquid level of the in the source wells were found to be critical for effective duckweed picking. Also, the destination wells must be pre-loaded with liquid for the release of the duckweed to happen, which can be done by the OT-2 just prior to duckweed loading. Hence, the entire experiment can be set up with the OT-2, streamlining and further accelerating experiment preparation.

The protocols to pick up the duckweed from a source well (on a 12-well plate) and drop off to a destination well (on a 96-well plate) were prepared using a web-based protocol designer tool provided by the manufacturer (Opentrons, 2022). Movements already defined in the protocol designer to handle liquids were employed for duckweed loading. Picking up duckweeds with an inoculation loop is based on capillary action between the hoops of the inoculation loop and the liquid surface, which causes the duckweed to climb on the loop and stick to it. We directed the inoculation loop to scan the source well surface with the “Touch tip” command, which passed the loop through where the duckweeds were most crowded in the well, and improved the success of duckweed picking. Dipping the inoculation loop into the destination well was enough to release the duckweed from the inoculation loop. A “Delay” command allowed the duckweed to drift away from the inoculation loop after dipping and before loop removal. A video of duckweed loading operation can be found in the SI. Figure 1 below illustrates the sequence of the duckweed loading protocol.

**Figure 1:**
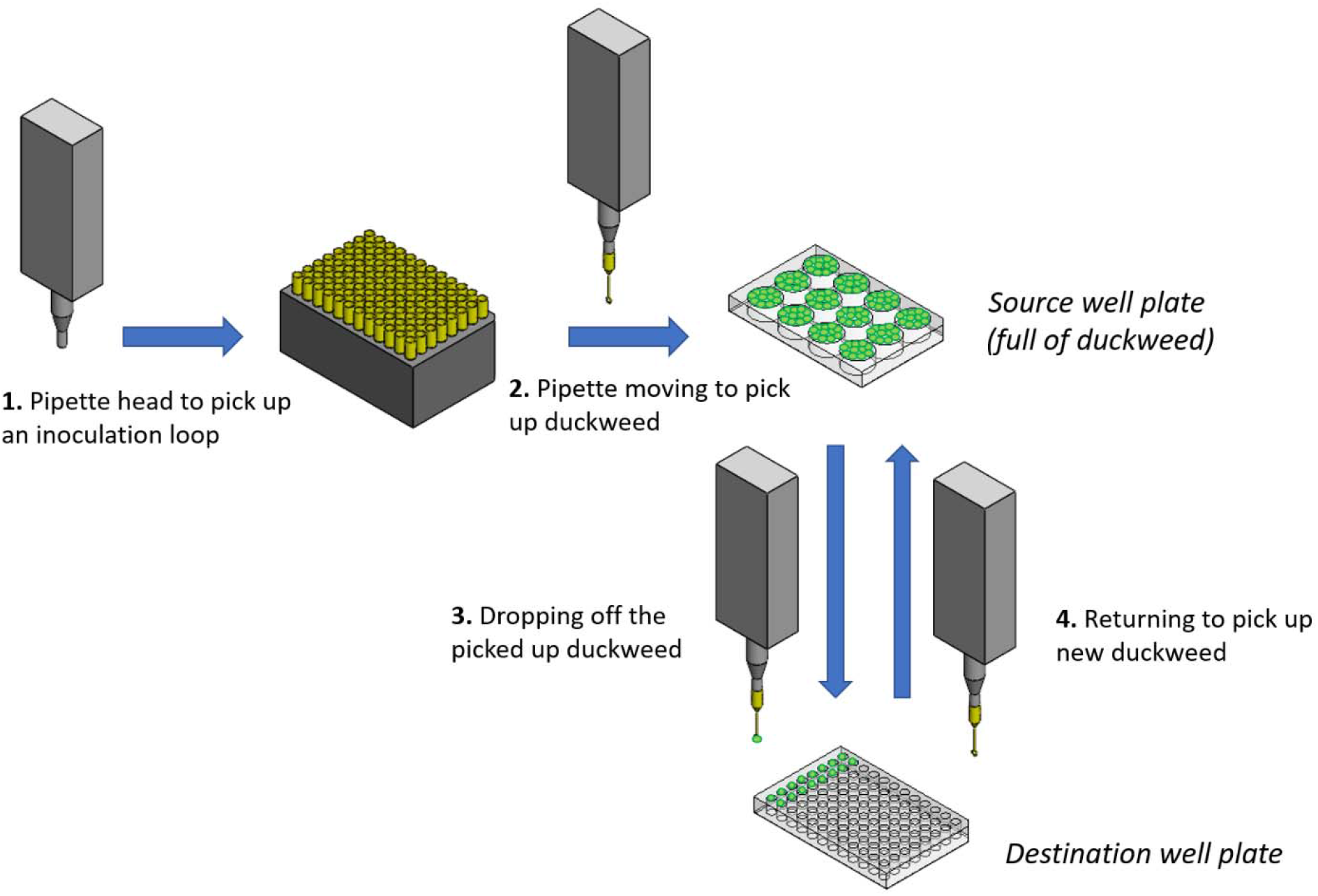
Schematics of the protocol sequence demonstrating the duckweed loading operation. This duckweed loading method operates in an open-loop fashion, i.e., no feedback mechanism is applied to this system. The open-loop method makes the system less complicated although it may fail to fill duckweed in some wells. The automated duckweed loading not only accelerates the entire experiment preparation process but also removes heavy-duty manual labor that otherwise has to be performed by a researcher. This manual labor had been the bottleneck to high-throughput experiment with duckweeds.

After loading the duckweed and other liquid experimental parameters using OT-2, the well plates were sealed with two membranes (Sigma Aldrich, Breathe Easy, Z380059, and Sigma-Aldrich, Breathe Easier, Z763624) to prevent cross-well microbial movement and to facilitate imaging, and moved to the growth chamber (see “Experiment Design”). The imaging of the samples can be also done autonomously and without interrupting the experiment, as discussed in the next subsection.

### Autonomous Imaging System

The imaging system consists of four high resolution (12.3 MP) cameras multiplexed over a Raspberry Pi 4 board, a camera holder, a linear actuator (Zaber Technologies, LC40B2000) and three transparent stages that hold the experimental well plates full of duckweeds. The entire system was built inside a growth chamber, which effectively also functions as a photo booth for duckweed samples. Figure 2 also presents the 3D model of the imaging system.

**Figure 2:**
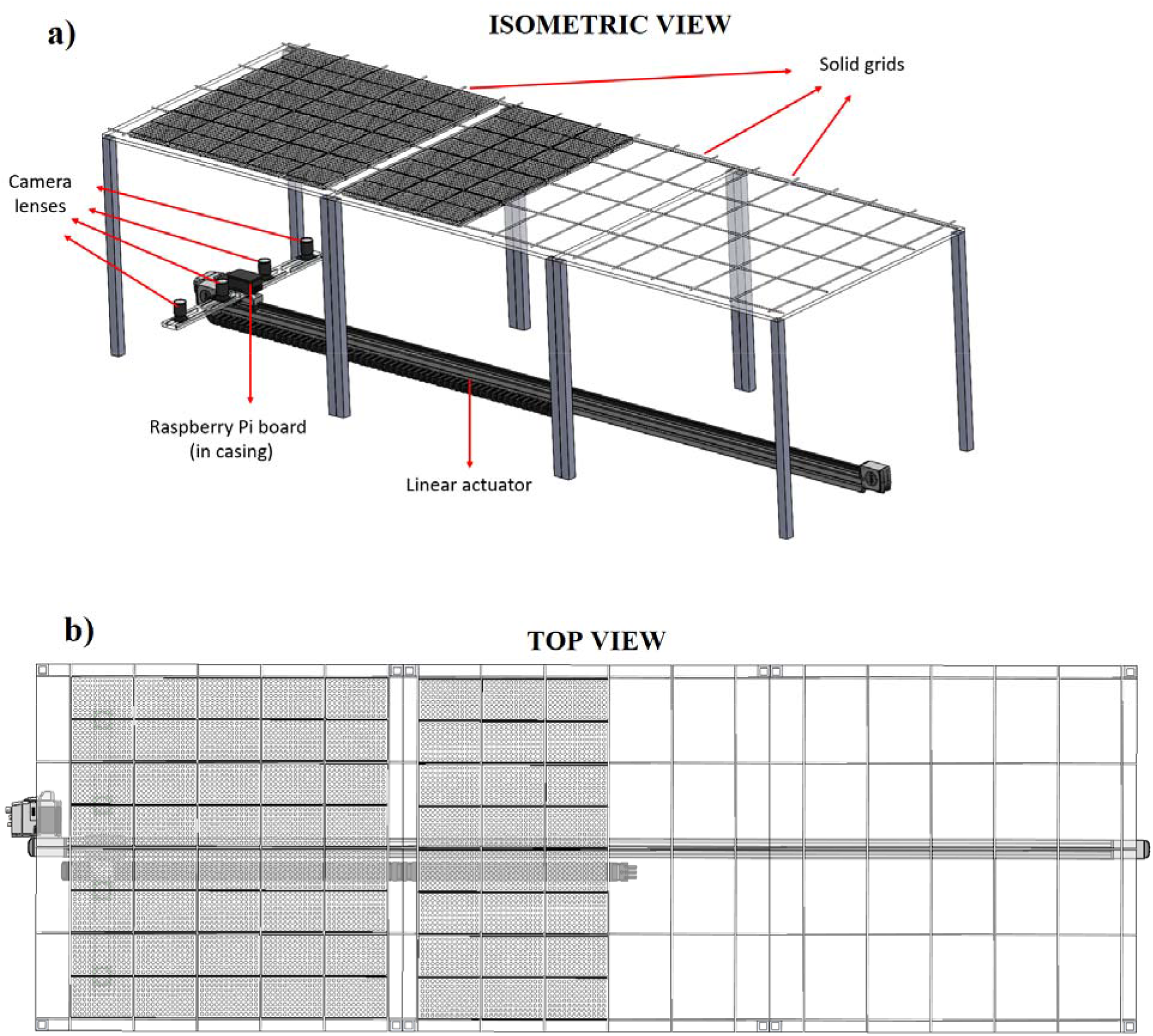
Autonomous imaging system built in the growth chamber a) in isometric view and b) in top view. The cameras take images from the bottom of the transparent stages on which the duckweed samples located. The solid grid on the transparent stages help locate the duckweed to prevent misalignment of samples. The linear actuator carries the imaging system to complete the imaging of all duckweed samples.

The cameras and the Raspberry Pi board were bolted on the linear actuator via the camera holder while the 96-well plates filled with duckweed samples were placed over the transparent stage. The position of the camera lenses can be adjusted to desired locations for each experiment by passing bolts through slots in the camera holder and the holes of the circuit boards (assembled with the camera lenses), sliding to horizontal position, and fixing with nuts. The transparent stages are made of 95% transparent acrylic that has a 75 cm x 75 cm footprint and 1 cm thickness. The linear actuator was located under the transparent stages enabling image capture from the bottom of the well plates in order to monitor the growth of the duckweed samples. We did not image the plates from above because the plate seals obscure images of the duckweeds.

When activated, the four cameras sequentially take photos of the well plates. Each camera captures an image covering two 96-well plates. Therefore, the well plates on the transparent stage are aligned such that there will be 8 well plates in each row. The linear actuator visits each row (total of 8 rows in this experiment) with the cameras and captures photos of all samples in under 10 minutes. Considering the physical limits of the growth chamber, the travel range of the actuator (2 m) and the size of a standard well plate, the imaging system is capable of imaging 120 well plates which means up to 11,520 experimental duckweed microcosms can be monitored in an experiment using 96-well plates.

To avoid detailed calibration before every experiment, positions of imaging system components are fixed inside the growth chamber. A solid mesh grid made of acetal rods was built on the transparent stages using epoxy and guarantees the well plates remain located in exact positions on the transparent stages. In a similar fashion, the positions of the transparent stages and the linear actuator were fixed with respect to the growth chamber walls.

All photos captured by the cameras are time-stamped and stored in Raspberry Pi memory, which are accessible through remote desktop connection for researchers in the network. Researchers can transfer the photos to their local computer for phenotype analysis using copy and paste. The control of the linear actuator and image capturing via Raspberry Pi cameras is automated over a Python script. The whole imaging system can be scheduled to operate autonomously at the desired time using this developed code. Hence, experimental imaging can be uninterrupted and is fully automated.

### Autonomous Phenotyping Tool

We created several graphical user interfaces (GUIs) to process the collected pictures, extract phenotypes, and visualize results using MATLAB Image Processing tools. Automatic image processing starts by extracting individual 96-well plate pictures from two well plates stored in each raw image, using the known geometries of the plates and growth chamber. Images are then further decomposed into individual images of each well on each well plate using computer vision. We implement the Hough transform to locate circles coinciding with the outline of each well and create a binary mask corresponding to the areas of each defined well. Not all wells can be determined using this approach due to noise and the difficulty in extracting the outline of a well. Therefore, we then superimpose a grid of circles corresponding to a 96 well plate onto the binary mask to predict the location of wells missed by the Hough transform. The high resolution of the raw images (4056 × 3040 pixels) means that each individual well image is 175×175 pixels, enough detail to observe changes in duckweed growth and morphology.

Analysis of duckweed growth is based on color thresholding. Here, we combined HSV-based and green intensity thresholding to predict whether a pixel is part of the duckweeds or is in the background. Color thresholding requires a uniformly white background to all wells, which our opaque plate seals achieve. The same settings were applied throughout the whole dataset. Our code converts the well image into a binary mask, which defines the footprint of duckweed from which the frond area and number of duckweeds per well can be extracted. Although the frond area and the number of duckweed fronds are the most significant parameters for this experiment, our code is also capable of examining the “greenness” of the duckweed. Greenness distinguishes unhealthy duckweeds from healthy ones, e.g., dying duckweeds turn white/yellow/grey while the healthy ones maintain their green color. Greenness in images of plants has also been associated with leaf nitrogen content, presumably because chlorophyll is both green and closely related to leaf nitrogen (Rorie et al., 2011; Schepers et al., 1992). Figure 3 illustrates the workflow of the phenotyping system.

**Figure 3:**
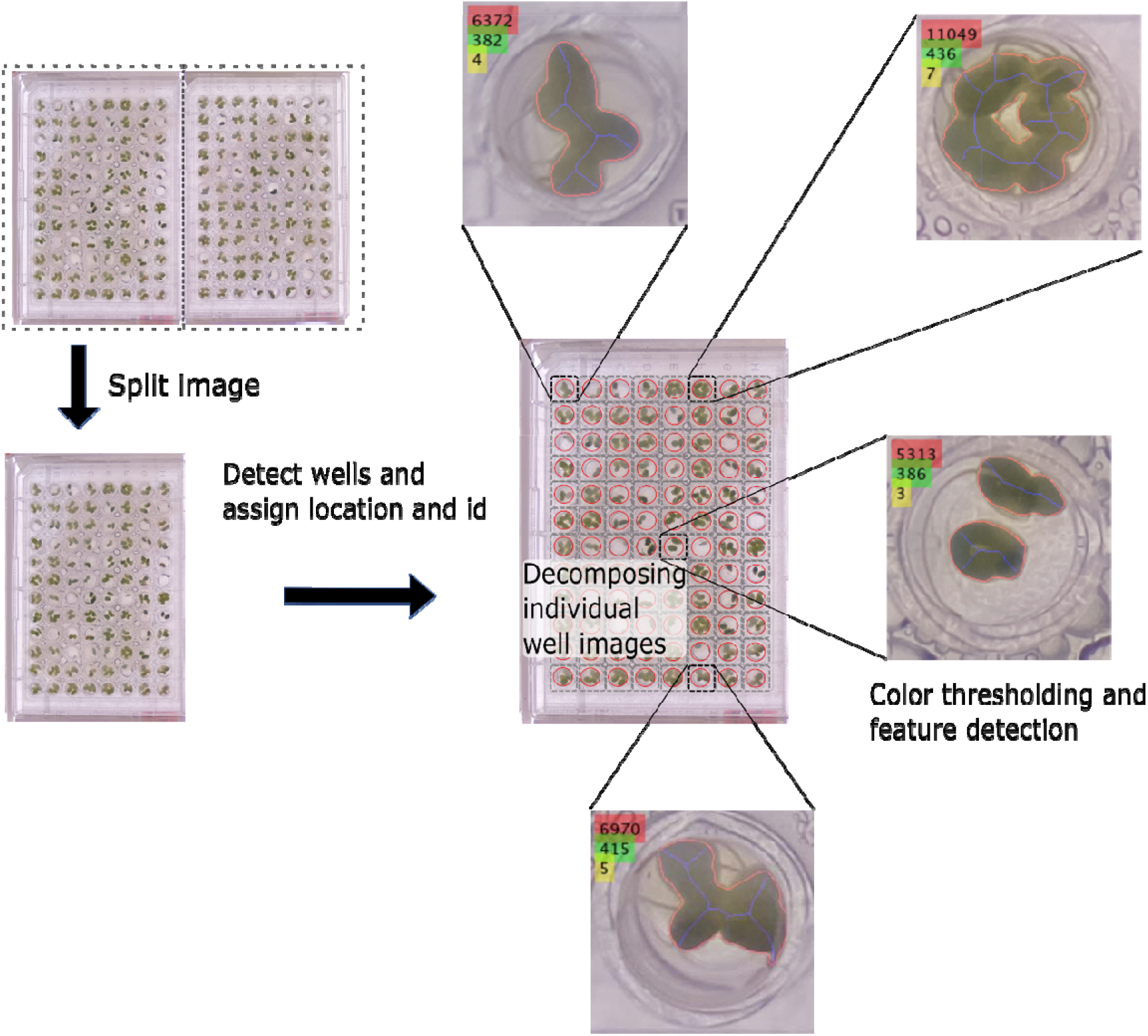
Workflow of the phenotyping tool. First, the raw image containing two well-plate images is split into individual well-plate images. Then, the wells are detected, labeled and decomposed into individual well images. Finally, color thresholding helps extracting the features of the duckweeds such as the total frond area, the greenness and the number of the fronds in a well.

We developed supplementary systems to handle post-processing hurdles resulting from slight deviations in experiment setup, such as poor well image extraction due to imprecise positioning of well plates (“tilted” images), or failing to detect the duckweed in the wells due to tone differences in color (lighting quality variation). The manual well detection interface, again created as a MATLAB GUI, helps researchers adjust automatic well detection results to correct any improper identification of the wells. Similarly, we created another GUI to allow researchers to adjust the automatic color thresholding to minimize the undetected duckweed fronds. This GUI allows the user to create a training sample, then uses a random search algorithm to improve color threshold by minimizing pixel misclassification. Figure 4 demonstrates the overview of the GUIs prepared. The details are discussed in detail in the SI.

**Figure 4:**
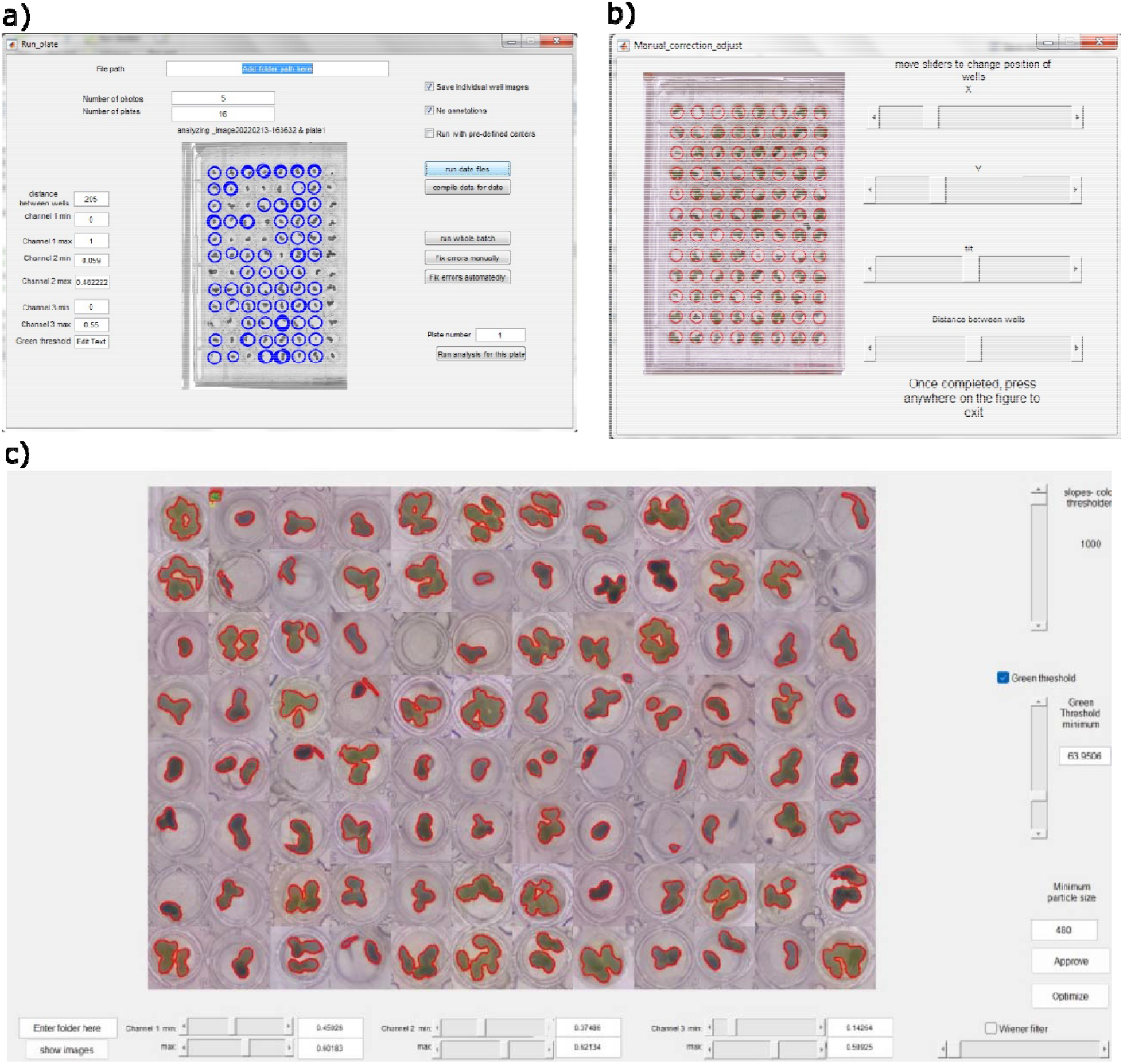
Graphical User Interface applications for automated phenotyping with main application to process image files, locate wells in each image and extract features from each well (a); manual correction tool allows user to manually fine tune the well location to ensure all duckweed area is accounted for in the analysis (b); color threshold settings optimization tool offers a visual way to change feature extraction settings and provides a tool to optimize settings using a training sample of images (c)

Phenotypic information gathered from image analysis included area of live fronds (calculated from pixel area using a conversion factor (pixels per well plate length ÷ length of well plate)^2^), frond number, and the greenness of fronds. Data analysis and visualization was carried out in R version 4.2.1 (R Core Team, 2022). Results were evaluated using linear mixed models to fit responses from the daily measurements of frond area and frond greenness. The models included fixed effects of microbes (presence/absence), N level, the number of days since the start of the experiment, and the interactive effect between microbes and N level. We fitted N level and the number of days as linear and second order polynomial terms. Random effects included well plate and whether the well was located on the edge or interior of the plate. Complete model results for the main and interactive effects of N, P, K, and microbes are available in (Wang et al., n.d.) (in preparation).

## Results

Our automated duckweed loading system was utilized to prepare an experiment with 6,000 microcosms of duckweeds growing in 2,000 nutrient and microbe treatments in 64 94-well plates. An OT-2 system first filled the experiment liquids using the system’s standard pipetting tools. Then, the OT-2 system outfitted with our new duckweed-picking tool filled the plates with experimental duckweeds. In duckweed loading, tests show up to 90% of experimental wells were successfully filled with duckweeds, with an average filling rate of 70%. In each well that was successfully filled, the automated system consistently loaded one single unit. The whole protocol to fill one 96-well plate takes 13 minutes. The 10-30% empty wells were manually filled by the researcher prior to the sealing of the well plates.

Our autonomous imaging system took photos of all units every day at 9 pm for 10 days and stored the raw photos in its memory (see Methods). The raw files were then extracted using remote desktop connection to a local computer where the autonomous phenotyping tool ran its post-processing color thresholding algorithms.

To showcase the abilities of the system for HTE, here we examine the effects of nitrogen levels on duckweed growth with and without microbes in our experiment with 6,000 experimental units. An in-depth analysis of results can be found in (Wang et al., n.d.) (in preparation).

The system successfully tracked the continuous growth of plants through time (Figure 5, Table 1), which would have been challenging and labor-intensive to do manually. Duckweeds grew quickly (large positive effect of Day: p < 0.001, Table 1). While the total frond area continued to increase over the course of the experiment, plant growth started to plateau around day 5 of the experiment (Figure 5, negative effect of Day^2^: p < 0.001). This may be explained by frond death occurring later in the experiment, or restricted frond growth within well boundaries as the frond area approached the total area of the well.

**Table 1:**
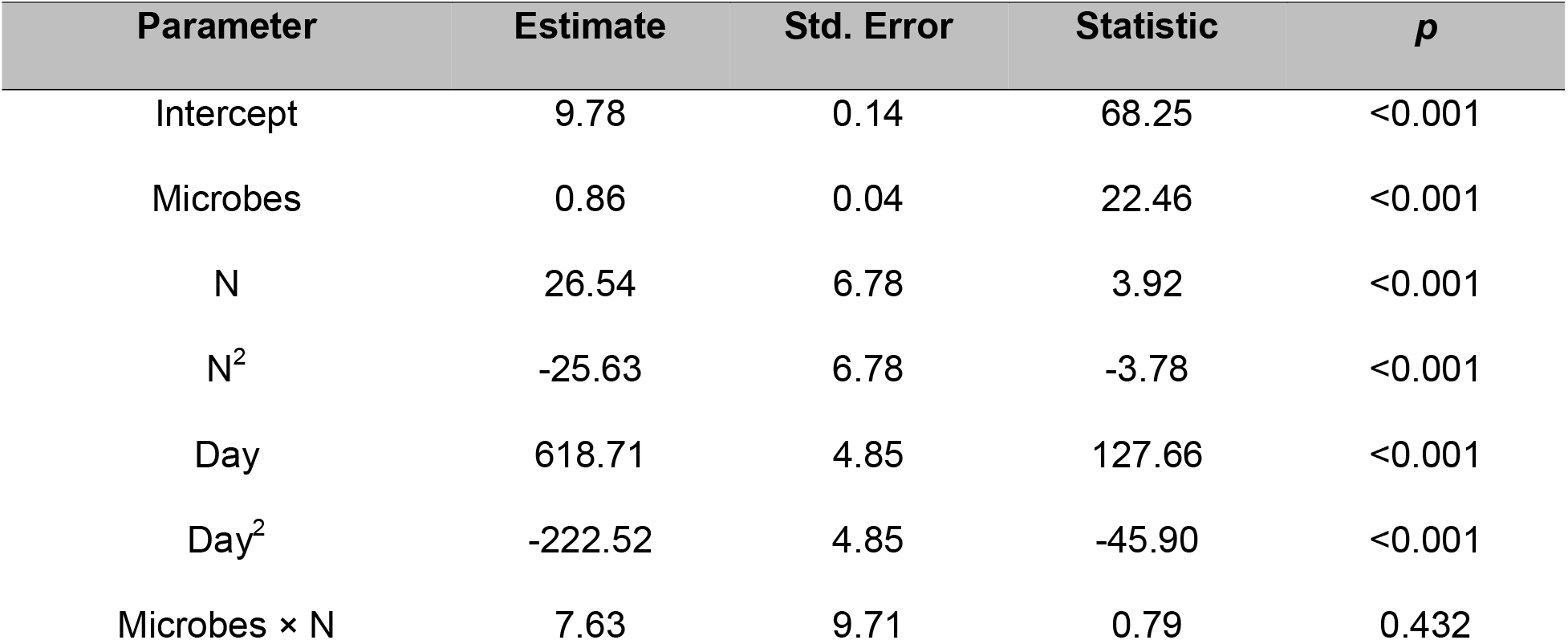
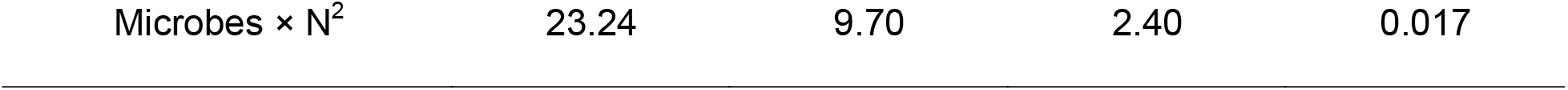
Model results for frond area. Values are reported for the fixed effects of each parameter, with “×” indicating an interactive effect between parameters. The following parameters were fitted as linear and second order polynomial terms: N = NaNO_3_ mg/L; Day = number of days since the start of the experiment.

**Figure 5:**
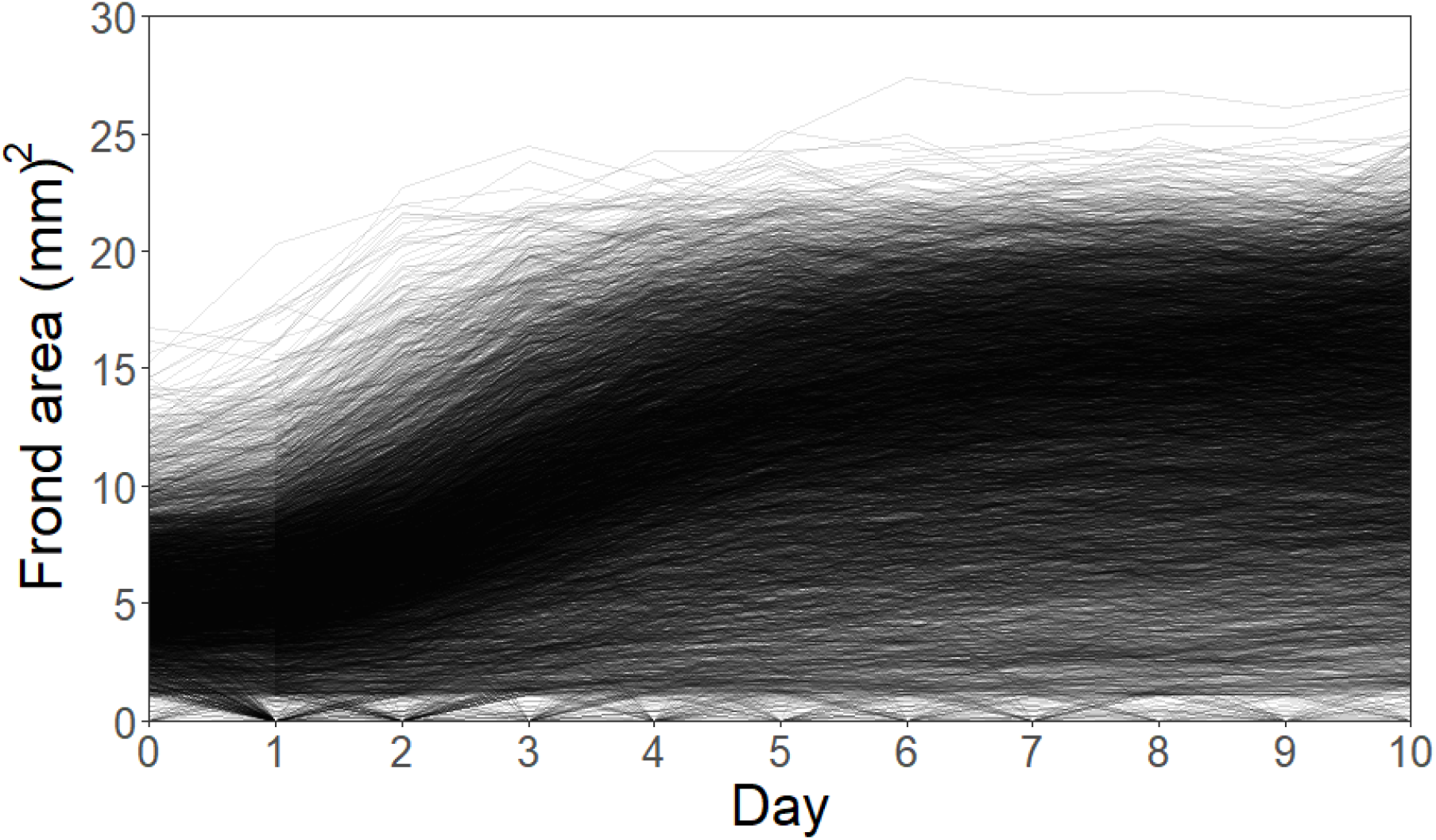
Growth of L. minor in all 6000 experimental units through time. Measurements of frond area were obtained from images taken once per day, over the 10 days of the experiment. Each line represents the plants located in one well.

In the absence of microbes and other nutrients, higher levels of N positively affected plant growth (positive N effect: *p* < 0.001), though not indefinitely, as revealed by a significantly negative quadratic effect (N^2^, *p* < 0.001), and reduced growth at the highest levels of N. The level of N that led to the most duckweed growth in the absence of microbes was 25 mg/L (Table 1, Figure 6). Plants inoculated with microbes experienced more growth overall (Microbes, p < 0.001, Figure 6). Inoculated plants also had the highest growth in midrange levels of N, but the addition of microbes shifted the level of nitrogen that supported the most duckweed growth, as there was a positive interactive effect between microbes and the square of nitrogen level (Microbes × N^2^: *p* < 0.017, but Microbes × N: *p* < 0.432, Table 1). In plants inoculated with microbes, the most growth occurred at 30 mg/L N, but growth did not decline as N increased beyond 30 mg/L (significant positive Microbes × N^2^ effect of similar magnitude to the negative N^2^ effect, Table 1, Figure 6).

**Figure 6:**
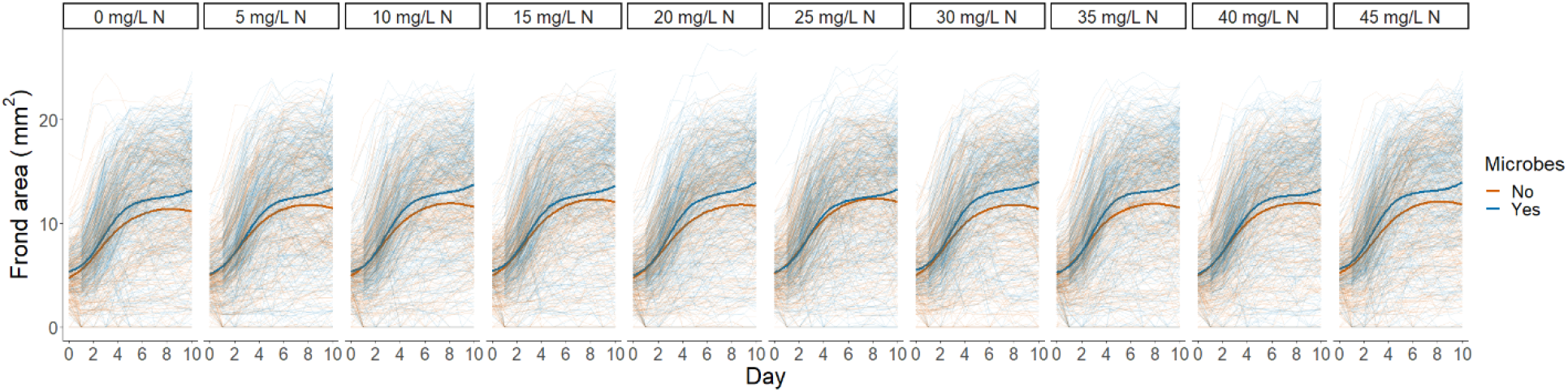
Growth of duckweeds across nitrogen levels. Measurements were obtained from images taken once per day for 10 days. Each line represents the area of fronds in one well across all other nutrient levels. Darker lines depict the smoothed conditional means of plants inoculated with microbes (blue) and plants not inoculated with microbes (orange).

The time-resolved data collected by the HTE system also revealed time-resolved effects on frond greenness, allowing a finer understanding of non-linearity in duckweed response to varying nitrogen levels and microbes. Frond greenness increased non-linearly with time but did not decrease by the end of the experiment (positive Day and Day^2^ effects, p < 0.001, Table 2). Instead, greenness briefly declined after loading into well plates, but then recovered, potentially due to the stress of moving from glass jar to the wells. The presence of microbes increased plant greenness (*p* < 0.001, Table 2), largely because plant greenness experienced a weaker initial decline when microbes were present. Unlike frond area, higher levels of N did not significantly affect plant greenness in the absence of microbes (N and N^2^ *p* > 0.1, Table 2). However, in inoculated plants, high levels of N resulted in marginally higher plant greenness (marginally significant Microbes × N^2^: *p* = 0.064, non-significant negative effect of Microbes × N: *p* > 0.1, Table 2)

**Table 2:** Model results for greenness of fronds. Values are reported for the fixed effects of each parameter, with “×” indicating an interactive effect between parameters. The following parameters were fitted as linear and second order polynomial terms: N = NaNO_3_ mg/L; Day = number of days since the start of the experiment.

| Parameter | Estimate | Std. Error | Statistic | p |
| --- | --- | --- | --- | --- |
| Intercept | 84.22 | 2.84 | 29.65 | <0.001 |
| Microbes | 3.46 | 0.28 | 12.15 | <0.001 |
| N | 33.09 | 50.24 | 0.66 | 0.510 |
| N<sup>2</sup> | -58.23 | 50.29 | -1.16 | 0.247 |
| Day | 2557.50 | 35.93 | 71.19 | <0.001 |
| Day<sup>2</sup> | 96.96 | 35.93 | 2.70 | 0.007 |
| Microbes × N | -39.35 | 71.94 | -0.55 | 0.584 |
| Microbes × N<sup>2</sup> | 133.05 | 71.92 | 1.85 | 0.064 |

## Discussion

The successful integration of our automated duckweed loading system, autonomous imaging system and autonomous phenotyping tool enabled a high-throughput duckweed experiment with 6,000 units.

Our automated duckweed loading system expedited the preparation process by removing the manual labor burden from the researcher. The current open-loop version based on the Opentrons liquid-handling robot OT-2 does most of the manual labor, and the success rate was was as high as 87%. The researcher was still needed to monitor the success of duckweed loading and to supplement the empty wells with duckweeds when necessary. Still, given the ease of implementation of the open-loop system, and amount of work required to improve the system performance marginally, we decided not to pursue closed-loop capability. Closed-loop capability could theoretically be realized by developing image-processing-based feedback mechanism to program the robot to re-attempt a failed duckweed transfer until it is achieved.

Our autonomous imaging system performs the imaging of the samples automatically, via remote control, and without disturbing the experimental conditions. One challenge we observed was the subtle vibration created by the environment support systems of the growth chamber, e.g., the fans working non-stop to maintain air flow in the chamber. This vibration caused the images to distort slightly during image capture, which may cause the frond area calculations to vary accordingly. The maximum variation of a frond area calculation (i.e. defined here as standard deviation for five images divided by the average area) is estimated to be ±4.7%. In order to eliminate any slight variation between captured images, we took five consecutive images using the same camera and averaged the phenotyping parameters in the post-processing.

Our autonomous phenotyping tool smoothly extracted phenotype and growth data in most cases. However, sometimes the color of the duckweed samples stayed near the color-thresholding limits our algorithm utilized (Figure 7). This resulted in “zero” frond area on one day, and a transition to a larger non-zero area on a later day with only a slight shift in frond color. Our optimization GUI helped tailor color thresholding limits to minimize these errors (as shown in Figure 7). We built a training set containing 250 images of wells pooled from different time points and treatments. For each training image, we varied the color threshold settings until obtaining a visually satisfactory detection of the duckweed area. Our optimization protocol then randomly searched for the set of HSV and green threshold settings that would minimize pixel classification error relative to all the training sample images. This optimization achieved a reduction in training error of ∼9%. Some of the remaining error can be in the training sample itself, as it may also contain human errors and bias. Most of the disparity is near the edges of the duckweed, which can be particularly difficult to define consistently with human or computer vision.

**Figure 7:**
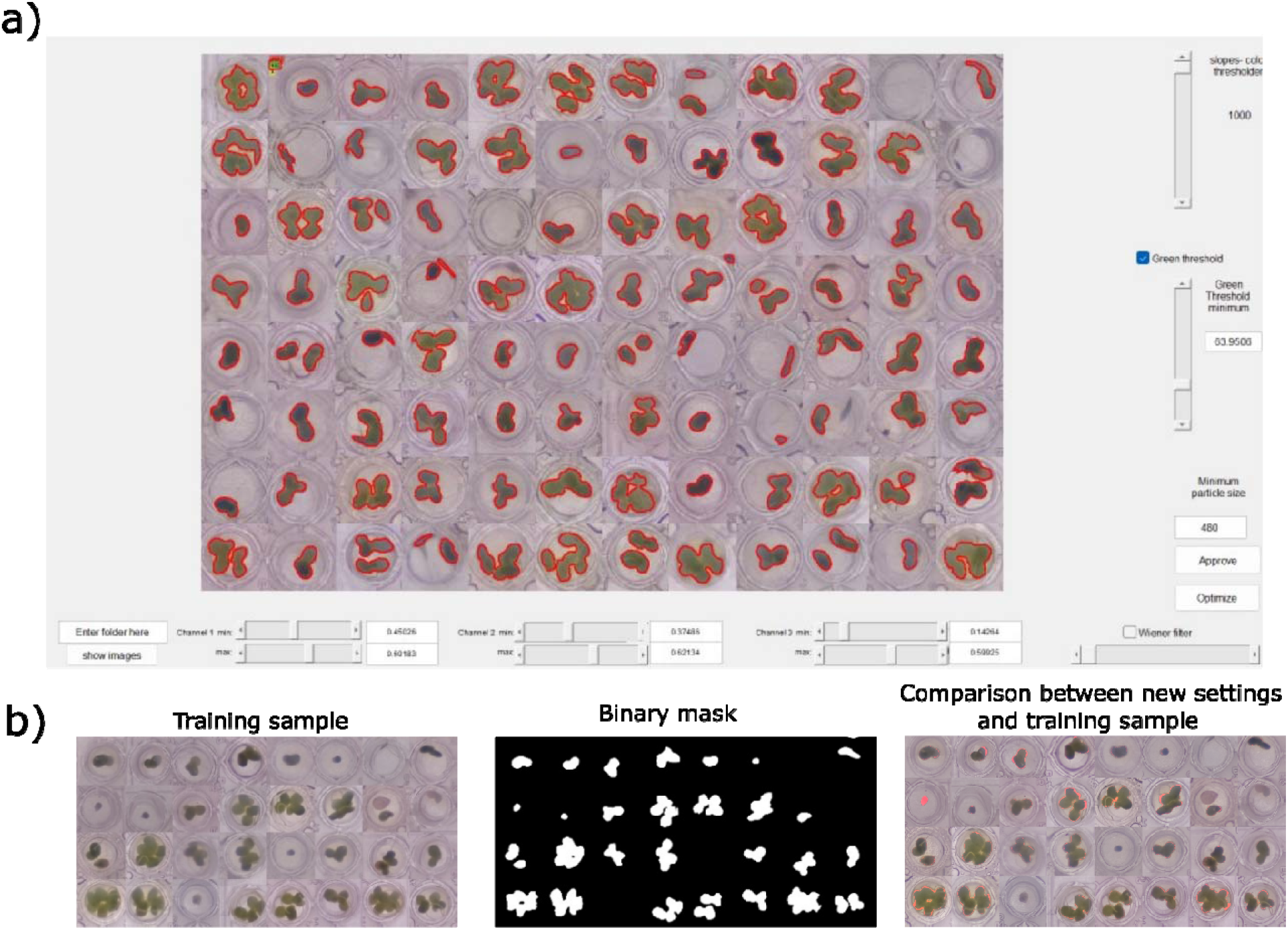
Optimization of duckweed detection. Here, we demonstrate our GUI to facilitate in the process of changing color threshold settings (a), followed by the training sample, binary mask for training sample, and a comparison between the performance of the optimal color threshold settings and the training sample, with areas of change in red (b).

Across our two biological response variables, high experimental throughput and automated phenotyping allowed a deeper understanding of the complex interactions between duckweeds and their environment. The time-resolved phenotype and growth data that our platforms provided across a high number of experimental treatments allowed finer-scale detection of time-resolved effects. Duckweed growth plateaued over time, and the nitrogen level that maximized growth depended on the presence or absence of microbes (Figures 5 & 6, Table 1). An initial decrease in plant greenness occurred only in treatments without microbes, and though plants in all treatments eventually increased in greenness, plants with microbes received an additional greenness boost, depending on nitrogen levels (Table 2)..

There are other interesting directions in which to develop high-throughput imaging systems for duckweed in the future. There are some studies (Zhang & Zhang, 2018) reporting 3D imaging of duckweeds to study microscopic features of duckweed, as well as fluorescence imaging (Oláh et al., 2021) and micro tomography (Jones et al., 2021). Perhaps most obviously, our automatic imaging and phenotyping systems could be extended to other organisms. Larger plants could be incorporated with automatic tissue-culture or seed-germination image-based assays. The automated imaging system is further capable of video recording, which could be applied to monitor mobile specimens. Automatic video monitoring and video processing might be desired for experiments with tiny invertebrates, such as *Daphnia, Artemia*, as well as small terrestrial insects. Indeed, video tracking phenotypes already exist for such organisms, and would only need to be integrated with our automatic imaging and automatic well-detection.

## Conclusion

Progress in the field of biology requires us to predictively decompose multiplicative effects of environmental and biological variation. These highly multi-factorial endeavors require increasingly vast quantities of data, which are prohibitively costly and time-intensive to generate manually. To sustain research in biology going forward, we need continued integration of biological model systems into automated platforms, like the one we outline here for duckweeds. There is particular value in flexible systems that can easily be adapted across organisms, such as our automated imaging and picking methods. Going forward, we expect that biology will make the most progress with improvements in automated handling of more different types of organisms and their requisite media, and an expanded imaging technology repertoire to integrate into automated phenotyping.

## References

Appenroth, K. J., Sowjanya Sree, K., Bog, M., Ecker, J., Seeliger, C., Böhm, V., Lorkowski, S., Sommer, K., Vetter, W., Tolzin-Banasch, K., Kirmse, R., Leiterer, M., Dawczynski, C., Liebisch, G., & Jahreis, G. (2018). Nutritional value of the duckweed species of the Genus Wolffia (Lemnaceae) as human food. Frontiers in Chemistry, 6(OCT). https://doi.org/10.3389/fchem.2018.00483

Birk, S., Chapman, D., Carvalho, L., Spears, B. M., Andersen, H. E., Argillier, C., Auer, S., Baattrup-Pedersen, A., Banin, L., Beklioğlu, M., Bondar-Kunze, E., Borja, A., Branco, P., Bucak, T., Buijse, A. D., Cardoso, A. C., Couture, R. M., Cremona, F., de Zwart, D., … Hering, D. (2020). Impacts of multiple stressors on freshwater biota across spatial scales and ecosystems. Nature Ecology and Evolution, 4(8), 1060–1068. https://doi.org/10.1038/s41559-020-1216-4

Caicedo, J. R., van der Steen, N. P., Arce, O., & Gijzen, H. J. (2000). Effect of total ammonia nitrogen concentration and pH on growth rates of duckweed (Spirodela polyrrhiza). Water Research, 34(15), 3829–3835. https://doi.org/10.1016/S0043-1354(00)00128-7

Cao, W., Zhou, J., Yuan, Y., Ye, H., Nguyen, H. T., Chen, J., & Zhou, J. (2019). Quantifying variation in soybean due to flood using a low-cost 3D imaging system. Sensors (Switzerland), 19(12), 1–12. https://doi.org/10.3390/s19122682

Choudhury, S. das, Samal, A., & Awada, T. (2019). Leveraging image analysis for high-throughput plant phenotyping. Frontiers in Plant Science, 10(April), 1–8. https://doi.org/10.3389/fpls.2019.00508

Cox, K. L., Manchego, J., Meyers, B. C., Czymmek, K. J., Harkess, A., & Czymmek, K. J. (2021). Automated imaging of duckweed growth and development. BioRxiv, 2021.07.21.453240. https://www.biorxiv.org/content/10.1101/2021.07.21.453240v1%0Ahttps://www.biorxiv.org/content/10.1101/2021.07.21.453240v1.abstract

David, A. S., Thapa-Magar, K. B., Menges, E. S., Searcy, C. A., & Afkhami, M. E. (2020). Do plant-microbe interactions support the Stress Gradient Hypothesis? https://doi.org/10.1002/ecy

Eyke, N. S., Koscher, B. A., & Jensen, K. F. (2021). Toward Machine Learning-Enhanced High-Throughput Experimentation. Trends in Chemistry, 3(2), 120–132. https://doi.org/10.1016/j.trechm.2020.12.001

Ge, X., Pereira, F. C., Mitteregger, M., Berry, D., Zhang, M., Hausmann, B., Zhang, J., Schintlmeister, A., Wagner, M., & Cheng, J.-X. (2022). SRS-FISH: A high-throughput platform linking microbiome metabolism to identity at the single-cell level. https://doi.org/10.1073/pnas

Gregori, F., Kapran, J., & Emilie, D. (2017). Filtration of Floating Particles by Collectors: Influence of the System Geometry on the Efficiency. Proceedings of the ASME 2017 International Mechanical Engineering Congress and Exposition, 1–7.

Jones, D. H., Atkinson, B. S., Ware, A., Sturrock, C. J., Bishopp, A., & Wells, D. M. (2021). Preparation, Scanning and Analysis of Duckweed Using X-Ray Computed Microtomography. Frontiers in Plant Science, 11(January), 1–17. https://doi.org/10.3389/fpls.2020.617830

Kehe, J., Kulesa, A., Ortiz, A., Ackerman, C. M., Thakku, S. G., Sellers, D., Kuehn, S., Gore, J., Friedman, J., & Blainey, P. C. (2019). Massively parallel screening of synthetic microbial communities. Proceedings of the National Academy of Sciences, 116(26), 12804–12809. https://doi.org/10.1073/pnas.1900102116

Krajncic, B., Slekovec-Golob, M., & Nemec, J. (1995). Promotion of Flowering by Mn-EDDHA in the Photoperiodically Neutral Plant Spirodela polyrrhiza (L.) Schleiden. In J. Plant Physiol (Vol. 147).

Lasfar, S., Monette, F., Millette, L., & Azzouz, A. (2007). Intrinsic growth rate: A new approach to evaluate the effects of temperature, photoperiod and phosphorus-nitrogen concentrations on duckweed growth under controlled eutrophication. Water Research, 41(11), 2333–2340. https://doi.org/10.1016/j.watres.2007.01.059

Newton, R. J. (1977). ABSCISIC ACID EFFECTS ON FRONDS AND ROOTS OF LEMNA MINOR L. American Journal of Botany, 64(1), 45–49. https://doi.org/https://doi.org/10.1002/j.1537-2197.1977.tb07603.x

Nguyen, B., Graham, P. J., Rochman, C. M., & Sinton, D. (2018). A Platform for High-Throughput Assessments of Environmental Multistressors. Advanced Science, 5(4), 1–9. https://doi.org/10.1002/advs.201700677

O’Brien, A. M., Yu, Z. H., Luo, D. ya, Laurich, J., Passeport, E., & Frederickson, M. E. (2020). Resilience to multiple stressors in an aquatic plant and its microbiome. American Journal of Botany, 107(2), 273–285. https://doi.org/10.1002/ajb2.1404

Oláh, V., Hepp, A., Irfan, M., & Mészáros, I. (2021). Chlorophyll fluorescence imaging-based duckweed phenotyping to assess acute phytotoxic effects. Plants, 10(12). https://doi.org/10.3390/plants10122763

Opentrons. (2022). Opentrons Protocol Designer BETA. https://designer.opentrons.com/

Rorie, R. L., Purcell, L. C., Mozaffari, M., Karcher, D. E., King, C. A., Marsh, M. C., & Longer, D. E. (2011). Association of “Greenness” in Corn with Yield and Leaf Nitrogen Concentration. Agronomy Journal, 103(2), 529–535. https://doi.org/https://doi.org/10.2134/agronj2010.0296

Schepers, J. S., Francis, D. D., Vigil, M., & Below, F. E. (1992). Comparison of corn leaf nitrogen concentration and chlorophyll meter readings. Communications in Soil Science and Plant Analysis, 23(17–20), 2173–2187. https://doi.org/10.1080/00103629209368733

Shears, N. T., & Ross, P. M. (2010). Toxic cascades: Multiple anthropogenic stressors have complex and unanticipated interactive effects on temperate reefs. Ecology Letters, 13(9), 1149–1159. https://doi.org/10.1111/j.1461-0248.2010.01512.x

Song, C., von Ahn, S., Rohr, R. P., & Saavedra, S. (2020). Towards a Probabilistic Understanding About the Context-Dependency of Species Interactions. In Trends in Ecology and Evolution (Vol. 35, Issue 5, pp. 384–396). Elsevier Ltd. https://doi.org/10.1016/j.tree.2019.12.011

Vasseur, F., Bresson, J., Wang, G., Schwab, R., & Weigel, D. (2018). Image-based methods for phenotyping growth dynamics and fitness components in Arabidopsis thaliana. Plant Methods, 14(1). https://doi.org/10.1186/s13007-018-0331-6

Wang, J., Kose, T., Lins, T., Pogoutse, O., Sinton, D., & Frederickson, M. E. (n.d.). Testing the outcome of host-microbiome interactions across 1000 environments. In Preparation.

Yang, W., Feng, H., Zhang, X., Zhang, J., Doonan, J. H., Batchelor, W. D., Xiong, L., & Yan, J. (2020). Crop Phenomics and High-Throughput Phenotyping: Past Decades, Current Challenges, and Future Perspectives. In Molecular Plant (Vol. 13, Issue 2, pp. 187–214). Cell Press. https://doi.org/10.1016/j.molp.2020.01.008

Yang, Y., Guo, Y., O’Brien, A. M., Lins, T. F., Rochman, C. M., & Sinton, D. (2020). Biological Responses to Climate Change and Nanoplastics Are Altered in Concert: Full-Factor Screening Reveals Effects of Multiple Stressors on Primary Producers. Environmental Science and Technology, 54(4), 2401–2410. https://doi.org/10.1021/acs.est.9b07040

Zhang, Y., & Zhang, N. (2018). Imaging technologies for plant high-throughput phenotyping: A review. Frontiers of Agricultural Science and Engineering, 5(4), 406–419. https://doi.org/10.15302/J-FASE-2018242

